# Probabilistic fine-mapping of transcriptome-wide association studies

**DOI:** 10.1101/236869

**Authors:** Nicholas Mancuso, Gleb Kichaev, Huwenbo Shi, Malika Freund, Alexander Gusev, Bogdan Pasaniuc

## Abstract

Transcriptome-wide association studies (TWAS) using predicted expression have identified thousands of genes whose locally-regulated expression is associated to complex traits and diseases. In this work, we show that linkage disequilibrium (LD) among SNPs induce significant gene-trait associations at non-causal genes as a function of the overlap between eQTL weights used in expression prediction. We introduce a probabilistic framework that models the induced correlation among TWAS signals to assign a probability for every gene in the risk region to explain the observed association signal while controlling for pleiotropic SNP effects and unmeasured causal expression. Importantly, our approach remains accurate when expression data for causal genes are not available in the causal tissue by leveraging expression prediction from other tissues. Our approach yields credible-sets of genes containing the causal gene at a nominal confidence level (e.g., 90%) that can be used to prioritize and select genes for functional assays. We illustrate our approach using an integrative analysis of lipids traits where our approach prioritizes genes with strong evidence for causality.

## Introduction

Transcriptome-wide association studies (TWAS) using predicted expression levels have been proposed as an approach to identify novel genomic risk regions and putative risk genes involved for complex traits and diseases.^1–3^ Since TWAS based on predicted expression only relies on the genetic component of expression, it can be viewed as a test for non-zero local genetic correlation between expression and trait.^1,4,5^ Significant genetic correlation is often interpreted as an estimate of the effect of SNPs on trait mediated by the gene of interest. However, this interpretation requires very strong assumptions that are likely violated in empirical data due to LD and/or pleiotropic SNP effects.^1–3,6–11^ Therefore TWAS has been mostly utilized as a test of association, in contrast to methods that attempt to directly estimate the mediated effect (i.e. Mendelian randomization^3,6–9^).

In this work, we show that the gene-trait association statistics from TWAS at a known risk region are correlated as a function of LD among SNPs and eQTL weights used in the prediction models. This effect is similar to LD-tagging in genome-wide association studies (GWAS) where LD within a region induces associations at tag SNPs (yielding the traditional Manhattan-style plots). Even in the simplest case where a single SNP causally impacts the expression of a gene which in turn causally impacts a trait, LD among SNPs used in the eQTL prediction models induce significant gene-trait associations at nearby non-causal genes in the region. The tagging effect is further exacerbated in the presence of multiple causal SNPs and genes. As an illustrative example, consider a risk region with 6 genes where a single SNP is causal for a single gene which impacts trait (causal gene in red; no other causal genes are present at this region, see Figure 1). Although genes 3 and 4 in Figure 1 have non-overlapping prediction weights due to different eQTL genetic regulation, LD among SNPs with non-zero prediction weights induce correlations in the TWAS statistics at genes 3 and 4. Estimating the correlation structure between predicted expression among nearby genes enables statistical fine-mapping over gene-trait associations. However, we note several confounding factors need to be addressed for unbiased inference. First, there is a body of evidence supporting horizontal pleiotropic effects from SNPs,^9,11,12^ which bias gene-trait association statistics and should be accounted for. Second, it is critical that TWAS fine-mapping approaches be robust if the causal mechanism is not steady-state levels of gene expression. Fine-mapping in these instances without controlling for confounding will result in a misspecified causal model and likely prioritize genes that tag causal mechanisms well.

**Figure 1:**
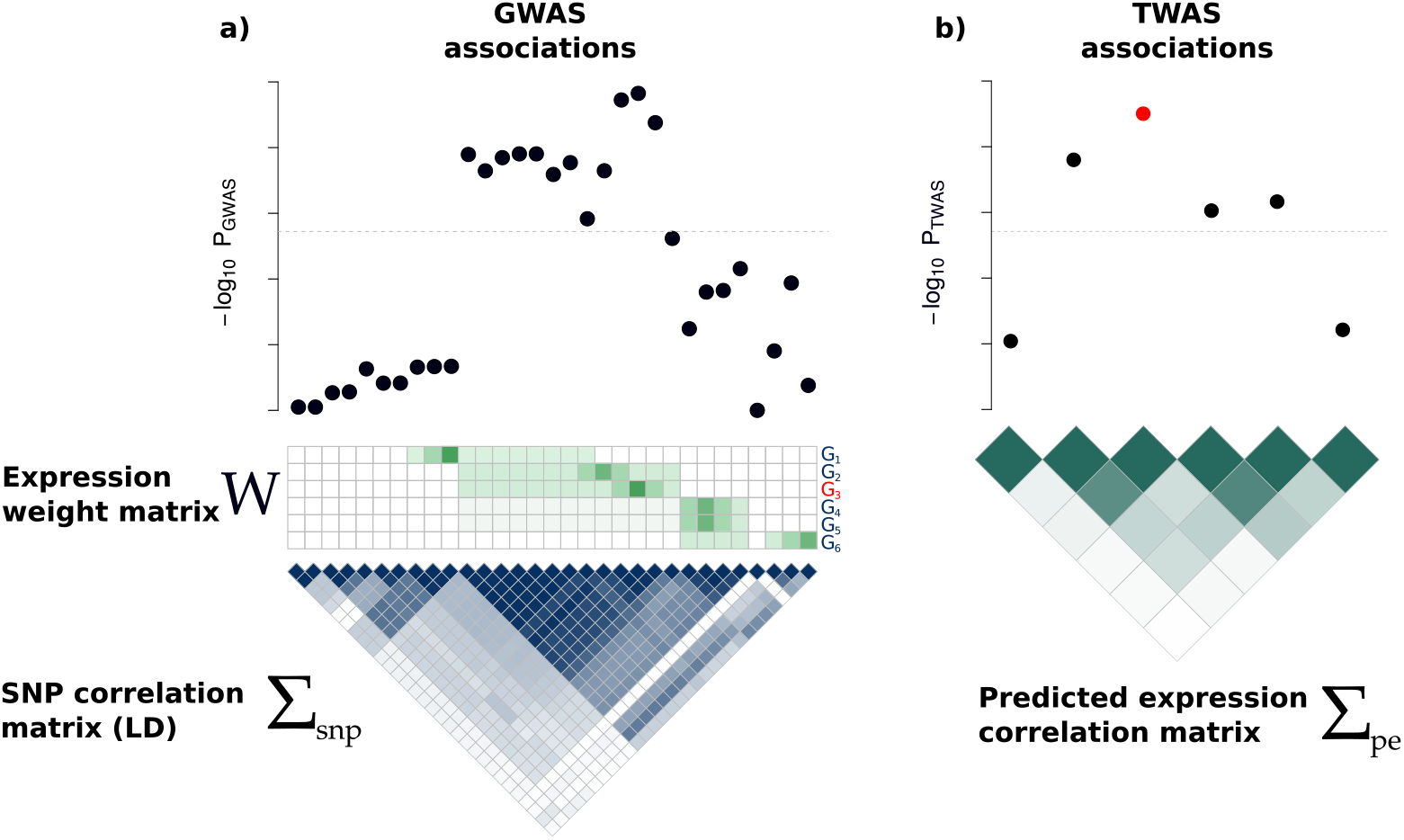
Illustration of the induced correlation structure for predicted expression. a) Top: Manhattan plot indicating strength of SNP association with trait. Middle: Expression weight matrix for 6 genes in the same region, with the causal gene in red. Each row corresponds to a gene and each column represents a SNP. Color indicates magnitude of eQTL effect. Bottom: The correlation structure (linkage disequilibrium, LD) across SNPs. Darker color indicates stronger correlation. b) Top: Transcriptome-wide association signal indicating strength of predicted expression association with trait. Bottom: Induced correlation of predicted expression. Darker color indicates stronger correlation between predicted expression levels. Dashed lines indicate the genome-wide (transcriptome-wide) significance threshold.

Here, we propose an approach to perform statistical fine-mapping over the gene-trait association signals while accounting for the correlation structure induced by LD and prediction weights used in the TWAS procedure and simultaneously controlling for certain pleiotropic effects. Our approach, FOCUS (Fine-mapping Of CaUsal gene Sets), takes as input GWAS summary data, expression prediction weights (as estimated from eQTL reference panels), and LD among all SNPs in the risk region, and estimates the probability for any given set of genes to explain the TWAS signal. Our approach extends probabilistic SNP fine-mapping approaches^13–15^ to estimate sets of genes that contain the “causal” genes (defined here as the gene responsible for the association signal) at a predefined confidence level (i.e. ρ gene credible set). FOCUS accounts for bias due to missing causal factors by including the *null model* as a possible explanatory factor in the credible set. We perform extensive simulations and show that FOCUS is unbiased in estimating the posterior probabilities and credible sets at a specified certainty when the causal gene is present in the data. When the causal tissue is unavailable and alternative tissues with correlated expression levels are used as a proxy, FOCUS maintains its performance under standard assumptions. FOCUS outputs posterior predictive checks^16^ of observed TWAS Z-scores to quantify model agreement given inferred posterior probabilities for causality. Finally, as a demonstration using real GWAS data, we apply FOCUS to four GWASs of lipids levels.^17^ We find that FOCUS prioritizes genes with established roles in LDL risk (e.g., *SORT1*).^18^

## Results

### Methods Overview

To disentangle between causal and tagging gene-trait associations at a TWAS significant region, we analytically derive the covariance structure among TWAS statistics as function of LD and eQTL weights used in prediction. Next, we model the entire vector of marginal TWAS association statistics (z_twas_) at all genes in a region (TWAS significant and not-significant) using a multivariate Gaussian distribution parameterized by the effect sizes at causal genes (**λ**_pe_), residual SNP-effects (**λ**_snp_), and the correlation structure induced by inferred expression weights (**Ω**) with LD (**V**) as

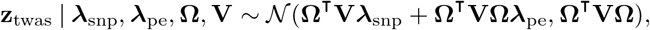

(see Methods). We control for bias resulting from pleiotropic effects of SNPs by including an intercept term that quantifies the average SNP effect sizes (**λ**_snp_) tagged by predicted expression (**Ω^Τ^Vλ_snp_**; see Methods). To allow for genes without prediction models in the relevant tissue (either due to QC and/or low power in eQTL studies), we leverage recent work demonstrating that eQTLs are largely shared across tissues^19^ and include prediction models from proxy tissues for such genes (see Methods). We employ a standard Bayesian approach to compute the marginal posterior inclusion probability (PIP) for each gene in the region to be causal. To avoid overfitting, we integrate out the unknown causal effects λ_pe_ using a multivariate Gaussian prior (see Methods). We use PIPs to compute *ρ*-credible gene-sets that contain the causal gene with probability p.^14^ To account for missing causal mechanisms either due to unpredicted expression or other latent functional mechanisms, we include the null model as a possible outcome in the credible set (see Methods). Lastly, we use a simulation-based procedure to compute posterior predictive checks^16^ that measure the FOCUS model’s goodness-of-fit given observed TWAS Z-scores.

### FOCUS prioritizes causal genes in causal-tissue simulations

To characterize the predicted expression correlation structure and to validate our framework, we used extensive simulations starting from real genotype data to generate expression reference panels and GWAS summary data (see Methods). We confirmed that non-causal genes in risk regions show significant association with trait as function of LD and eQTL weights (see Supplementary Figure 1), which motivates fine-mapping to prioritize genes causally impacting trait. We simulated complex trait under a variety of architectures to assess the performance of 90%-credible gene-sets computed using FOCUS (see Methods). When the causal gene was assayed in its relevant tissue, we found 90%-credible gene-sets contained 0.91 (S.D. 0.06) of causal genes across simulations on average (see Figure 2). We saw accuracy under general *ρ* was stable, with credible gene-sets being well-calibrated across various values of *ρ* (see Supplementary Figure 2). FOCUS models an intercept term to control for pleiotropic SNP effects (i.e. λ_snp_) tagged through predicted expression. In simulations where a fraction of SNPs directly impacted downstream trait, we computed credible sets after estimating an intercept term 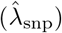 and found a decrease in performance (see Figure 2; see Methods). Next, we varied sample size across GWAS and reference eQTL datasets. Intuitively, we found improved performance for FOCUS to detect causal genes as sample size increased (see Figure 2, Supplementary Figure 3). Sample size for the eQTL reference panel affected performance to a larger degree than GWAS sample size, consistent with earlier reports.^1^ For example, at *N*_eQTL_ = 100, we found 90%-credible gene-sets contained the causal gene in 88% of simulations, which is significantly fewer when compared with 94% for *N*_eQTL_ = 500 (Mann-Whitney-U P = 5.46 × 10^−9^). Next, we explored how underlying heritability of expression at causal genes impacts prioritization. Heritability defines the prediction upper bound for SNP-based approaches,^20,21^ and we expect performance to improve as non-zero heritability is easier to detect. We confirmed that performance increased with heritability of causal gene expression (see Figure 2). For example, we simulated gene expression having heritability 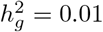 and inferred inferred eQTL weights using *N*_eQTL_ = 500 and found a significant decrease in performance (Mann-Whitney-U *P* < 2.2 × 10^−16^). Similarly, we looked at the role of the prior effect-size distribution for predicted gene expression^4^ and found performance to remain relatively stable for a wide range of values (see Supplementary Figures 4, 5).

**Figure 2:**
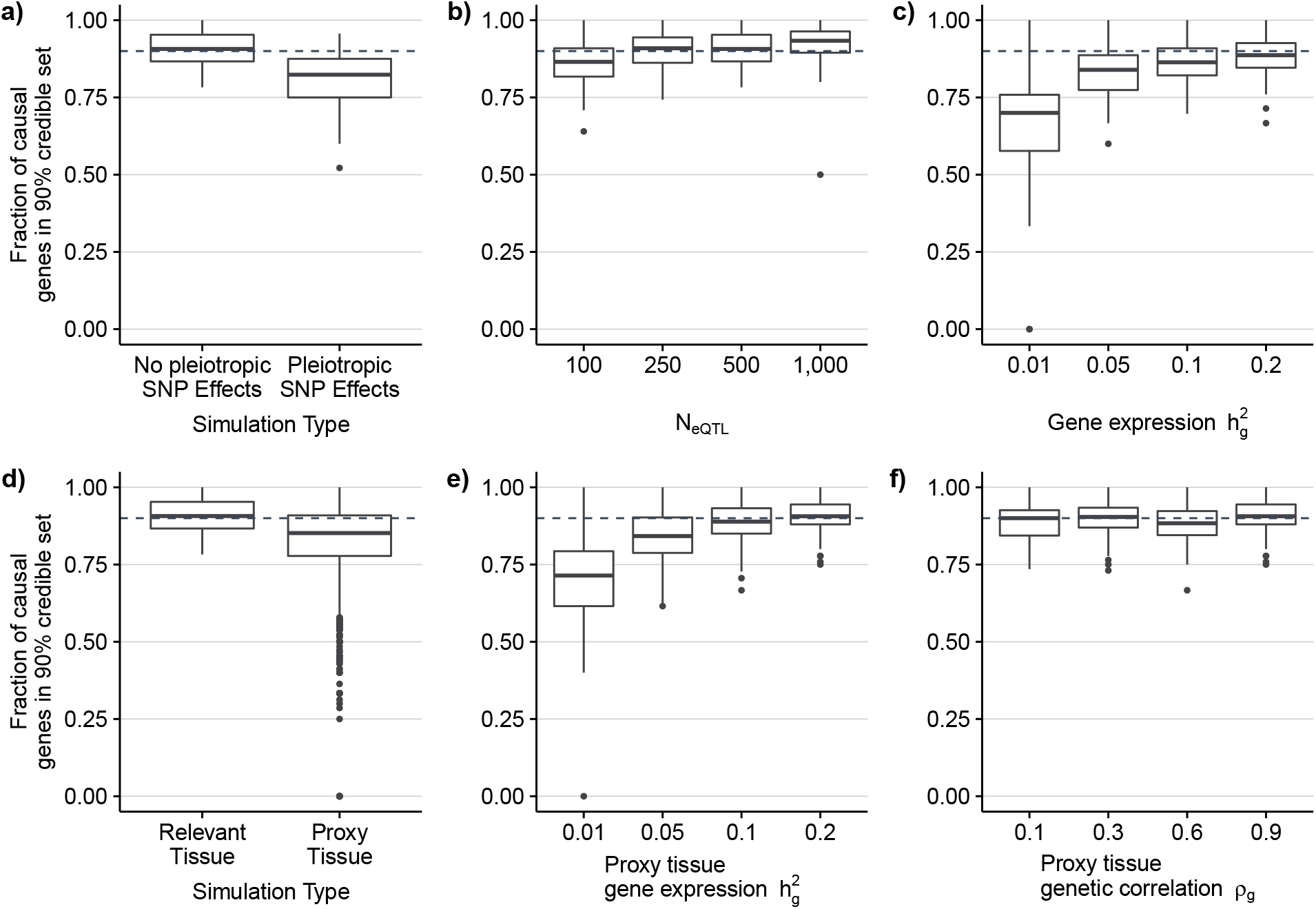
Credible gene-sets are well-calibrated in simulations. Box-plots represent the distribution of the fraction of causal genes captured in the 90% credible set over simulations (see Methods). a) Simulations with and without pleiotropic SNP effects on trait. Prediction models were trained using the relevant (i.e. causal) tissue. b) Calibration as a function of eQTL reference panel sample size. c) Calibration as a function of heritability of causal gene expression. d) Calibration using prediction models trained using proxy tissue measurements. e) Calibration using proxy tissue when heritability of reference gene expression varies compared with fixed 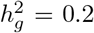 in the relevant tissue. f) Calibration using proxy tissue when genetic correlation of reference gene expression and gene expression in the relevant tissue varies.

### FOCUS remains stable when using proxy tissues

Next, we investigated the performance of FOCUS when the causal gene in the relevant tissue is missing, but is measured in a different tissue (see Methods). In real data a gene may act through a tissue that is difficult to assay in large sample sizes, but may have similar cis-regulatory patterns in tissues that are easier to collect (e.g., blood, LCLs). Indeed, several studies^1,4,19,22^ established cis-regulated gene expression levels exhibit high genetic correlation across tissues and functional architectures. The intuition in this approach is that the loss in power from using the correlated tissue is offset by the gain in power due to larger sample size. To simulate gene expression in a proxy tissue, we drew correlated effect sizes at the same eQTLs for the gene expression reference panel (see Methods). Here, we consider a causal gene to be successfully fine-mapped if its corresponding proxy tissue model is in the 90%-credible gene-set. When sample size for eQTL in the relevant- and proxy-tissues are the same, but heritability in proxy tissue is lower than the relevant-tissue, we found a significant loss in accuracy, with 90% credible sets capturing the causal gene 0.83 (S.D. 0.08) of simulations compared with 0.91 (S.D. 0.06) when averaging over values of *ρ_g_*. (Mann-Whitney-U *P* = 3.4 × 10^−13^; see Figure 2). This effect was not observed when heritability of proxy tissue gene expression was at least that of expression in the relevant-tissue (Mann-Whitney-U *P* = 0.06). For example, when expression in the relevant tissue was 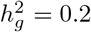, but 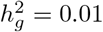 in the proxy, we found 90%-credible gene-sets contained the causal gene in significantly fewer simulations (0.79 versus 0.89; Mann-Whitney-U *P* < 2.2 × 10^−16^), which suggests that when causal eQTLs are shared across tissues, increased heritability of expression increases power to detect the causal gene. Surprisingly, we found correlation of effect-sizes at shared eQTLs to play no significant role in performance when heritability of expression in the relevant and proxy tissue is kept the same (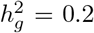; see Figure 2, Supplementary Figure 6). In the case of zero correlation between effect sizes at the same eQTL SNPs, this result can be interpreted as pleiotropic effects on independent molecular traits, which are known to be difficult to differentiate from a causal effect.^1,3,9^ Collectively, these results demonstrate that FOCUS is relatively robust to model perturbations and performs well when underlying tissue-specific causal genes are represented by proxy tissue eQTL weights.

### FOCUS is robust to missing causal molecular mechanisms

Our model predicts that predicted expression of nearby genes will be correlated due to linkage between eQTL SNPs, which results in correlated test statistics among gene-trait associations. If predicted expression for the causal gene is not included, nearby genes will likely be prioritized in fine-mapping. This scenario is analogous to absent causal SNPs under SNP fine-mapping approaches. FOCUS controls for this scenario through two mechanisms (see Methods). First, FOCUS explicitly models the null (i.e. no gene-trait relationship for all nearby genes) as a possible outcome when computing credible gene-sets. We tested the performance of FOCUS in simulations when there is no relationship between expression and trait, and found the null model was contained in the 90%-credible gene-set in 0.98 of our simulations, indicating that FOCUS is accurate under the null. We next performed experiments using simulations where causal gene expression effects downstream trait, but has been pruned from the data before testing. We found the null model in 0.59 (S.D. 0.08) of 90%-credible gene-sets (see Figure 3), which was a significantly greater percentage compared with simulations where the causal gene was present (0.14, S.D. 0.08; Mann-Whitney-U *P* < 2.2 × 10^−16^). Second, FOCUS estimates a single intercept term at each region to account for pleiotropic SNP effects on trait. When the predicted causal mechanisms are absent from the data, we expect the intercept to increase in magnitude, as SNP effects mediated through missing mechanisms will be indistinguishable from pleiotropic effects. Indeed, we found an enrichment for significantly non-zero intercept estimates (i.e. 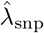) when the causal mechanism was absent compared with complete-data simulations (Fisher’s exact P = 2.9 × 10^−3^). Altogether, we find FOCUS is robust in the challenging setting of prioritizing the null model when causal expression is missing.

**Figure 3:**
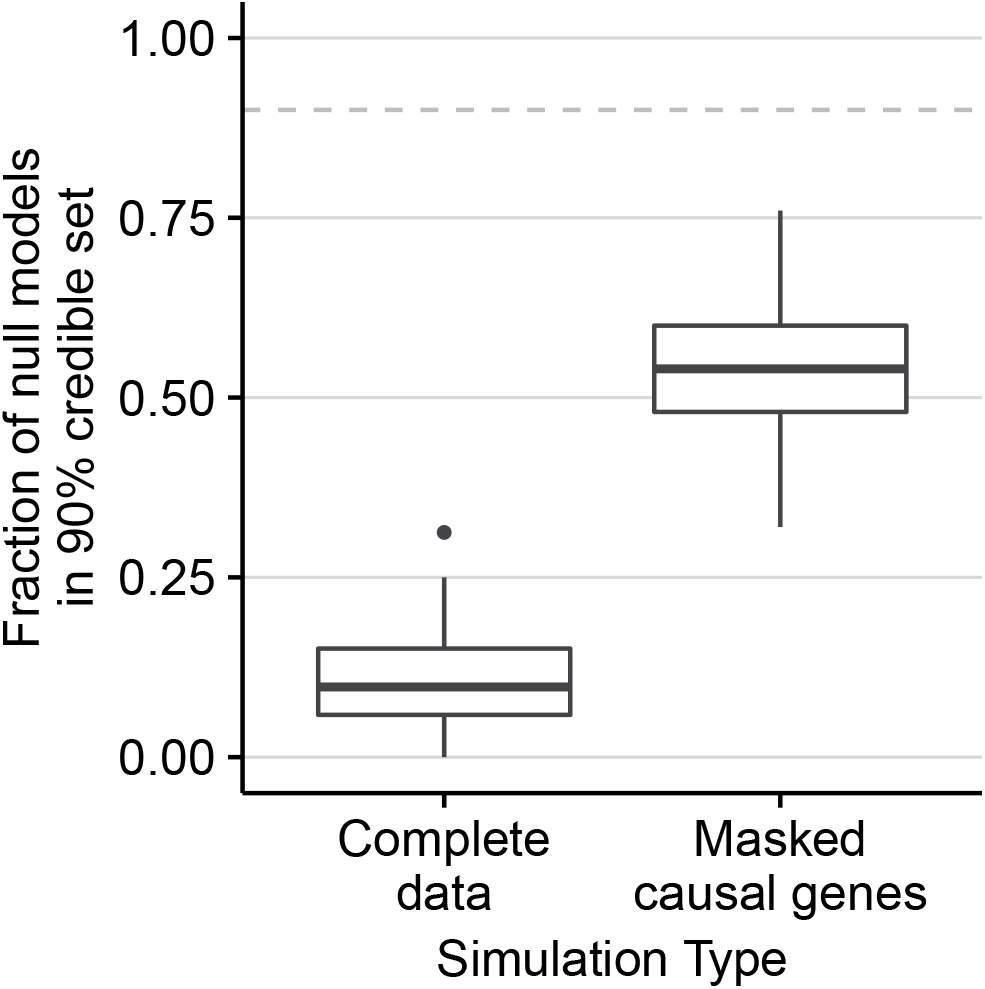
FOCUS credible-sets alleviate bias when the causal gene is missing. “Masked” indicates simulations where the causal genes are pruned before analysis. For comparison, we include results under the complete-data simulation pipeline where causal genes are tested. Box-plots represent the distribution of the proportion of null models captured by 90%-credible gene-sets in simulations.

### FOCUS improves resolution for fine mapping causal genes

Having demonstrated that FOCUS computes well-calibrated credible gene-sets under a wide range of parameters we next sought to quantify the resolution of credible gene-sets to identify causal genes. In particular, we estimated the average number of genes captured in the credible set. We found 90%-credible gene-sets contained 4.4 genes on average (S.D. 1.9) in the relevant-tissue simulations, which resulted in an average 47% of predicted genes per risk region (see Figure 4). We found a similar number of genes in 90%-credible gene-sets across simulations when varying model parameters and sample sizes (see Supplementary Figures 7-12). While we advocate the use of credible-sets in practice rather than thresholding on PIPs, for completeness we prioritized genes using PIPs for direct comparison with TWAS p-values (see Methods). We found prioritizing genes using PIPs outperformed TWAS p-value ranking at capturing underlying causal genes (see Figure 4). For example, at a false positive rate of 5%, FOCUS identifies 162% more causal genes than a simple rank of marginal TWAS statistics. A unique feature of FOCUS is that it allows for multiple causal genes at a given region; in this scenario FOCUS attains a a gain of 213% more causal genes compared to that of 132% for single causal regions (see Supplementary Table 2). Overall, FOCUS accurately prioritizes causal genes from non-causal genes with the largest gains when multiple causal genes exist at risk regions.

**Figure 4:**
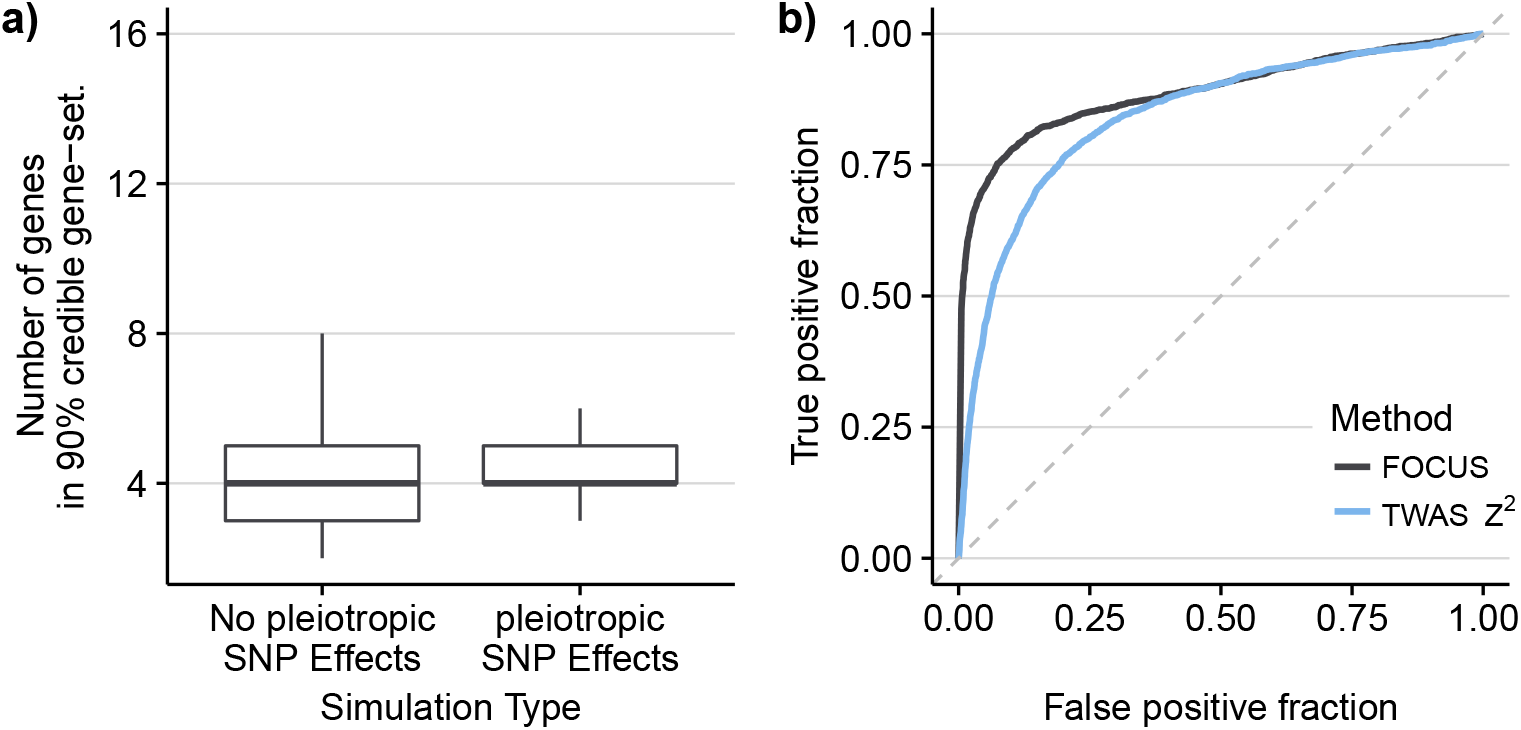
FOCUS accurately prioritizes causal genes in simulations. Box-plots represent the distribution of the total number of causal genes captured in the 90% credible sets over simulations (see Methods). a) Simulations with and without pleiotropic SNP effects on trait. Prediction models were trained using the relevant (i.e. causal) tissue. b) ROC curve computed using PIPs versus TWAS *Z*^2^ for each gene.

### Application to lipids GWAS data

Having validated our fine-mapping approach in simulations, we illustrate FOCUS by re-analyzing a large-scale GWAS of lipids measurements^17^ with eQTL weights from adipose tissue. We assume the relevant tissue for expression driving lipids is adipose given its well-characterized role.^23–26^ To account for missing gene prediction models, we incorporate gene expression models for genes not predictable at current sample sizes from adipose tissue across 45 tissues measured in 47 reference panels. In detail, for a gene without a predicted model in adipose tissue, we include the prediction model with best accuracy across all other tissues (see Supplementary Table 1; see Methods). Of the 26,292 known genes in RefSeq (ver 65),^27^ we found 12,663 covered in our data with the remaining 2,614 genes not found in RefSeq. Adipose-prioritized TWAS identifies 301 (202 unique) significant genes at 108 (63 unique) independent regions after accounting for the total number of per-trait tests performed (*P* < 0.05/15,277; see Supplementary Figures 13-15; Table 1; Supplementary Table 3). Of the 160 (89 unique) risk regions found through GWAS, 75 (46 unique) overlapped significant TWAS results, which is increased compared with earlier work^29^ that found 25% overlap between GWAS and eQTL at risk regions (see Table 1).

**Table 1:** Summary of gene-based fine-mapping in lipids GWAS risk regions. A GWAS risk region is defined to be a LD-block defined by LDetect^28^ harboring at least one genome-wide significant SNP (*P* < 5 × 10^−8^) reported in ref.^17^ A TWAS gene is a gene whose predicted expression reaches transcriptome-wide significance of *P* < 0.05/15, 277.

| Lipid Trait | GWAS risk regions | GWAS risk regions with any TWAS genes | TWAS genes at risk regions | Genes in 90%-credible sets |
| --- | --- | --- | --- | --- |
| High-density lipoprotein (HDL) | 43 | 18 | 64 | 30 |
| Low-density lipoprotein (LDL) | 36 | 20 | 56 | 40 |
| Total cholesterol (TC) | 51 | 24 | 73 | 53 |
| Triglycerides (TG) | 30 | 13 | 33 | 25 |
| Overall (unique) | 160 (89) | 75 (46) | 226 (146) | 148 (100) |

Having performed a TWAS across lipids traits, we next sought to prioritize putative causal genes using our framework. We applied FOCUS at the 75 GWAS risk regions with evidence for regulatory action on genes driving lipids levels to compute PIPs and estimate credible sets of genes at each of the regions (see Methods). We found that observed risk regions can be explained by 1.5 causal genes on average, with 61/75 risk regions containing fewer than 2 causal genes in expectation (see Supplementary Figure 18). The average maximum PIP across credible sets was 88% (and decreased exponentially for lower ranked genes; see Supplementary Figure 17). Together, these results imply that most risk regions can be explained by a single causal gene. Using computed PIPs, we estimated 90%-credible gene-sets for each risk region and found a significant reduction in the number of prioritized genes (mean 1.9), compared with transcriptome-wide significant genes (mean 3.2; one-sided Mann-Whitney-U P = 7.24 × 10^−4^; Supplementary Figure 17; Supplementary Table 4). As a positive control, we examined the 1p13 locus for LDL, as this region harbors risk SNP rs12740374 which has been shown to perturb transcription of *SORT1* and impact downstream LDL levels.^18^ We found 4/34 genes included in the 90% credible set, of which *SORT1* had a posterior probability 95% (see Figure 5).

**Figure 5:**
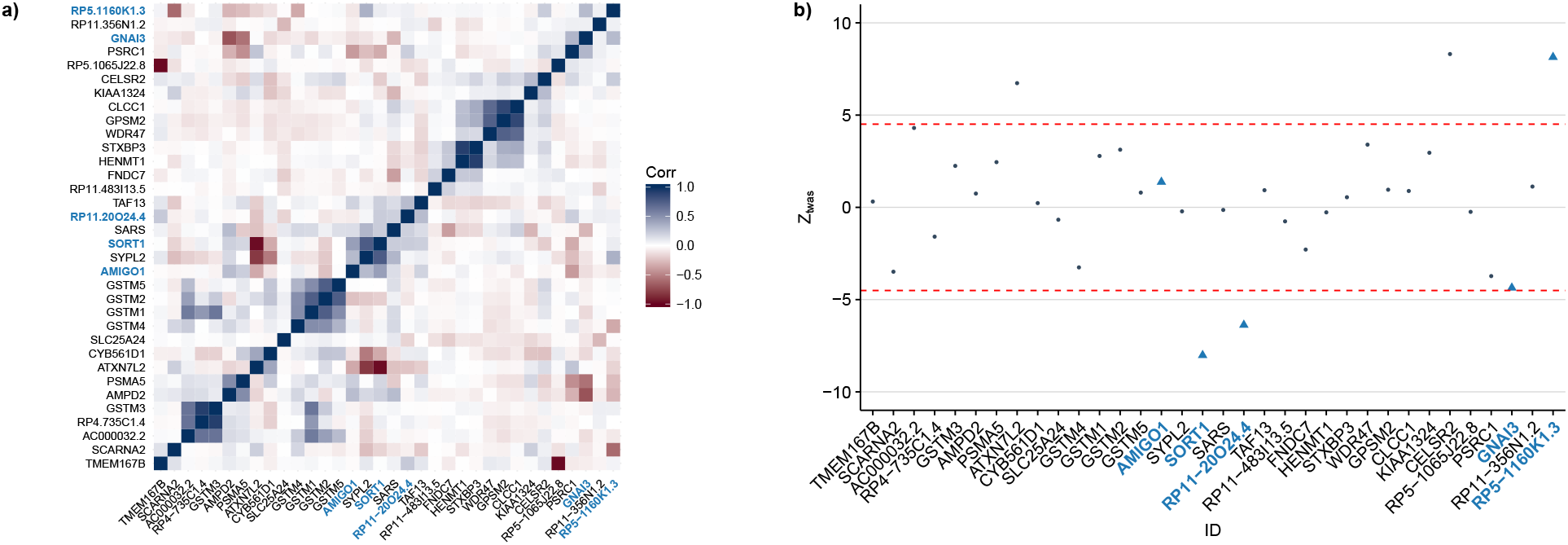
1p13 locus for LDL. a) Correlation for predicted expression at 1p13 locus. Genes in the 90%-credible set are labeled in light blue. b) TWAS Z-scores at 1p13 locus. Each point represents the association strength for each tested gene. Genes in the 90%-credible gene-set are labeled in light blue. Dashed red lines indicate transcriptome-wide significance threshold.

Next, we investigated regions whose 90%-credible gene-sets contained the null model (i.e. regions with weaker evidence for models of gene expression driving risk). An instance that contains the null model in its credible set may be partially consistent with observed association between expression levels and trait being due to chance. We found 25/75 instances of the null model captured in credible sets for lipids traits (see Supplementary Table 4), which suggests most overlapping GWAS risk regions are more consistent with risk contributed from cis-regulated expression levels, compared with statistical noise explaining observed signal. PIPs output by FOCUS are conditioned on the FOCUS model being correct. If FOCUS’s model does not accurately capture the underlying generative process then PIPs will be biased. We used a simulation procedure (see Methods) to quantify model fit for each gene and found the FOCUS model largely agreed ı with observed data (i.e. TWAS Z-scores; see Supplementary Figure 19).

## Discussion

In this work we presented FOCUS, a fine-mapping approach that estimates credible sets of causal genes ı using prediction eQTL weights, LD, and GWAS summary statistics. We demonstrated FOCUS adequately controls false positives in null simulations and outperforms straightforward p-value ranking in identifying causal genes when genes at a region impact downstream trait. We found 90%-credible gene-sets to be largely stable across a variety of simulations, with the biggest impact in performance due to eQTL reference panel sample size and SNP-heritability of gene expression. We applied FOCUS to four lipids TWASs (e.g., HDL, LDL, triglyceride, and total cholesterol levels) and found *SORT1* correctly identified as a putative causal ı gene. Interestingly, our real-data results in lipids suggests most regions can be explained by a single causal gene. Overall, our results highlight the utility of using credible sets in prioritizing causal genes by jointly assigning posterior probabilities, that are both easily interpretable and comparable across genes and regions.

In addition to providing a quantification of the confidence in how many genes need to be validated to identify ı the causal genes in the region, our probabilistic approach yields several benefits. First, FOCUS naturally allows for multiple causal SNPs and genes while integrating gene-effect sizes using conjugate priors; this is particularly important as recent works have shown that allelic heterogeneity (i.e. multiple causal genes i and SNPs at a region) is pervasive in both eQTL and GWAS.^19, 30^ Second, in this work, we investigate predicted gene expression, but FOCUS could generally be applied to other predicted molecular traits with ¦ an established role in complex trait etiology (e.g., alternatively spliced exons^31, 32^). For example, several recent works have supporting evidence for splice variation playing an important role in driving risk of schizophrenia.^33, 34^

We showed our approach is well calibrated under various simulations and robust to perturbations in model assumptions; however, several limitations still exist. First, our model assumes that complex trait or disease ı risk is a linear function of steady-state expression levels at causal genes. Several works have demonstrated that risk prediction using a linear combination of predicted steady-state or observed expression levels can outperform standard SNP-based models,^33, 35^ which supports a linear model of gene expression impacting complex trait or disease risk. However, higher-order models that capture complex regulatory networks of transcription factors and gene expression may also reflect underlying biology. As reference gene expression data sets grow in size, accurately modeling these assumptions may be possible. Similarly, if risk is mediated through context-specific expression and not steady-state expression levels, then FOCUS will have a loss in performance. Second, while our simulations used GBLUP^36, 37^ throughout for its straightforward implementation, we recommend a cross-validation approach to select the best fitting linear model (e.g., GBLUP, BSLMM^38^) using the ratio of out-of-sample prediction accuracy normalized by the total SNP-heritability ı of gene expression, which is implemented in the FUSION framework.^1,33^ Third, when the causal gene is untyped in the data, our approach will partially inflate posterior probabilities at tagging genes. We attempt to mitigate this scenario by adding an intercept term to the model and incorporating gene models measured in proxy tissues. We caution that our simulated results using proxy-tissues were performed using a model where causal eQTLs are shared between proxy- and relevant tissues and have correlated effect sizes, which is equivalent to a random-effects model. This assumption may be violated in real data if causal eQTLs are tissue-specific. Recent work, however, has demonstrated that a large number of eQTLs are indeed shared across tissues.^19^ Fourth, we took a tissue-prioritizing approach by preferentially using eQTL weights in adipose tissue given its known role in lipids^23–26^ for our real-data analysis. This approach may not always be possible for complex traits or diseases with less understood biology. However, recent work has shown that the most relevant (i.e. likely causal) tissue for complex traits can be accurately estimated using eQTL data.^39^ Coupled with estimation of causal tissue, we suggest prioritizing genes with high normalized prediction accuracy in related tissues. We note that our results were strongly dependent on sample size in the eQTL reference panel, which is reflected in expression prediction accuracy. We therefore, recommend prioritizing eQTL data with sample sizes greater than 100 if possible and performing inference on genes with robustly non-zero SNP-heritability. Despite our modeling assumptions and limitations, our approach is a step towards accurately prioritizing gene-sets.

## Online Methods

### Notation

We denote scalar variables with italicized lower-case letters (e.g., *z*). Vectors are denoted with bold lowercase letters (e.g., **z**). Scalar entries for a vector are indexed with a subscript (e.g., *j*th element of **z** is *z_j_*). We denote matrices with bold capital letters (e.g., **X**, its transpose **X^Τ^**) and index rows with a subscript (e.g., **X***_j_*). We indicate *L* block-column partitions for matrix (vector) **X** as **X** = (**X**^(1)^, …, **X**^(L)^).

### Model and sampling distribution of marginal TWAS summary statistics

We model quantitative trait for *n* individuals y by a linear combination of expression levels for *m* genes 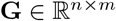 as

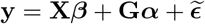

where 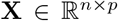 is the centered and variance-standardized genome-wide genotype matrix at *p* SNPs, *β* are the *p* pleiotropic effects of *X* on **y**, *α* is the vector of causal effects for the m genes and 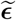 is random environmental noise with 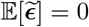 and 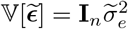 We extend our definition by also defining **G** as a linear function of underlying genotype and environment, which is governed by **G** = **XW** + **E**, where 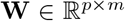 is the eQTL effect-size matrix, and 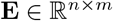 is environmental noise. Our updated model is

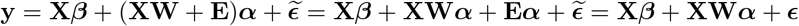

where 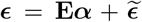 is the total contribution from environment, which we parameterize as 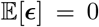 and 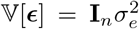 which is valid provided independence of errors holds. If causal eQTL effect-sizes W were known, we could prioritize putative susceptibility genes by estimating α using regression. Unfortunately, effect-sizes **W** are unknown and must be estimated from data (e.g., BSLMM,^38^ GBLUP^36,37^). Because inferring eQTL effect-sizes genome-wide is challenging, models typically focus only on *cis*- or local-SNPs at each gene. Let predicted expression be defined as **Ĝ** = **XΩ** when **Ω** is estimated from data. Our local-SNP model for *L* independent genetic regions is given by,

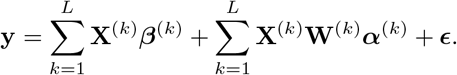

Here we describe the sampling distribution of marginal TWAS Z-scores obtained from an association test. For simplicity, we focus our attention to genes in a single genomic region and drop the (·) notation. Specifically, we compute the marginal association *z_j_* of gene *j* with **y** through a transcriptome-wide association study as,

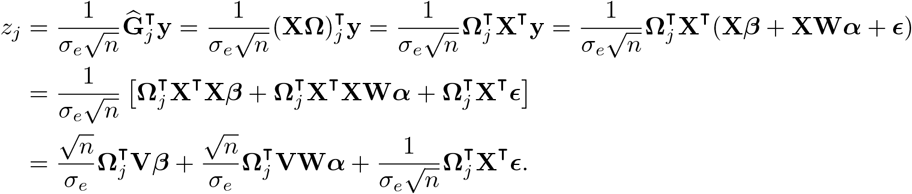

where **V** = *n*^−1^**X^Τ^X** is the SNP correlation (LD) matrix. The marginal association statistics for m nearby genes are determined by,

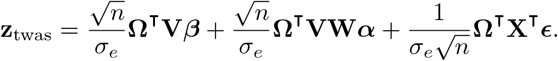

Assuming weights **Ω** and causal gene effects **α** are fixed, we can compute the expectation and variance of the association statistics as,

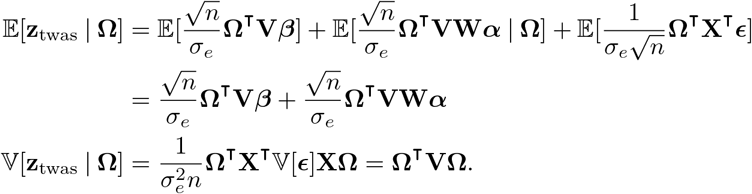

To simplify notation we re-parameterize the causal effects as a non-centrality parameter (NCP) at the causal I genes by 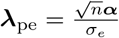. We note that 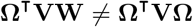, but given large-enough sample sizes we expect **Ω^Τ^VΩ** to approach **Ω^Τ^VW**^1^. We denote predicted expression covariance as 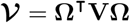. The NCP λ_pe_ governs the statistical power of rejecting the null of no effect of predicted expression on trait (*α* = 0). We parameterize *β* similarly as 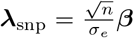. If we assume 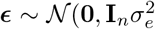, then our sampling distribution for **z**_twas_ is given by,

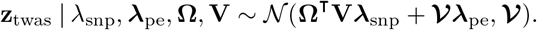

This formulation asserts that observed marginal TWAS Z-scores are the linear combination of NCPs at causal genes convoluted through the covariance structure of predicted expression 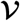 and tagged pleiotropic effects from SNPs **Ω^Τ^V***β*. Likewise, the resulting covariance structure 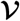 is the the product of the underlying LD structure of SNPs **V** and the weight matrix learned from expression data **Ω**.

Computing the likelihood of **z**_twas_ as described requires knowing 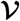, λ_snp_, and λ_pe_, which are unknown a-priori. First, we can estimate 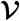 using available reference LD panels (e.g., 1000 Genomes^40^) and inferred expression weights **Ω**. Second, while we can estimate *β* from data, it will typically be the case that *p* ≫ *m*, which limits inference. To account for this, we make the simplifying assumption that λ_snp_ = 1_*p*_λ_snp_ when conditioned on 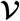 and λ_pe_, which is similar to methods in robust Mendelian Randomization.^9,11,12^ Third, estimating λ_pe_ directly from data is also likely to overfit. To bypass this issue, we treat λ_pe_ as a nuisance parameter and assume that 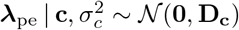 where 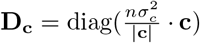 is the scaled prior causal effect variance and **c** is a binary vector indicating if ith gene is causal. Incorporating this prior for causal NCPs enables us to integrate out λpe, which results in the variance component model,

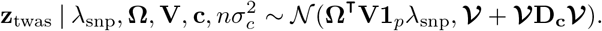

Under this model the variance in **z**_twas_ is due to uncertainty from finite sample size 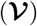 as well as uncertainty in the underlying causal NCPs 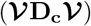. In principle, we can estimate 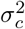 using Empirical Bayes; however, this comes at a significant computation cost, as estimation would need to be performed for each causal configuration c across risk regions. To mitigate this hindrance, we set 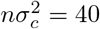, which is similar to what we observe at transcriptome-wide significant regions.

Equipped with our likelihood model for **z**_twas_, we take a Bayesian approach similar to fine-mapping methods in GWAS to compute the posterior distribution of our causal genes **c**,

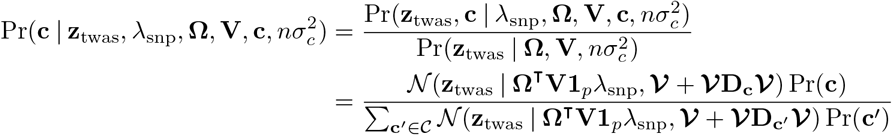

where 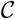 is the set of all binary strings of length *m*. We assume a Bernoulli prior for each causal indicator **c**_*i*_ ~ Bernoulli(*p*). In practice, we set *p* = 1 × 10^−3^. This assumption is likely violated when signal for **z**_twas_ is low, and we recommend only including regions with at least one transcriptome-wide significant gene. We compute the marginal posterior inclusion probability (PIP) for the ith gene as

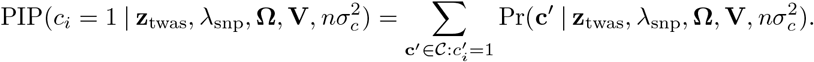

Alternatively, we can compute PIPs using Bayes factors for each model (see Supplementary Note). PIPs offer a flexible mechanism to generate gene-sets for functional followup. We use an approximate approach that takes the top *k′* genes until a percentage *ρ* of the normalized-posterior mass is explained.

### Model validation using the posterior predictive distribution

To test the validity of the FOCUS model at GWAS risk regions in real data, we use a posterior predictive sampling procedure.^16^ This approach alternates between sampling causal configurations c from the posterior distribution and sampling Z-scores 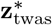 from the generative distribution after conditioning on the causal configuration. This enables us to compare the distribution of simulated data with our observed statistics **z**_twas_. When our observed data **z**_twas_ are not fit within reasonable bounds of the simulated data we can be more confident that the FOCUS model and computed PIPs are inconsistent with the actual data generating process. Specifically, at each risk region with m genes we perform the following:

For trial *t* ∈ [*T*]

1. For gene *i* ∈ [*m*]

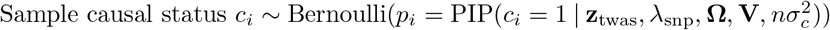
2. Sample simulated Z-scores 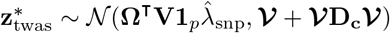
3. Output 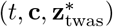

We compute a posterior Z-score (and p-value) of model fit for the *i*th gene as 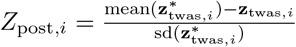.

### Simulations

We simulated TWAS association statistics starting from real genotype data and gene definitions. To simulate genotype samples, we first partitioned genotype data for 489 individuals of European ancestry in 1000Genomes^40^ into independent LD blocks as defined by LDetect.^28^ We annotated LD-blocks with all genes in RefSeq^27^ whose transcription start site was flanked by region boundaries. To simulate GWAS and expression reference panel genotypes we sampled standardized genotypes using the multivariate Gaussian approximation 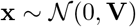 where **V** is LD estimated from the 1000Genomes samples. For both GWAS panel and eQTL reference panel, we simulated gene expression of each gene in the LD-block annotation list by selecting 1 or 2 causal SNPs preferentially located near 100kb of the TSS and then computed **G** = **Xw+ϵ** where **X** is the *n×p* centered and standardized genotype matrix, **w** are the causal eQTL effects, and 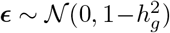 is random environmental noise. To simulate expression in two correlated tissues, we sample eQTL effects at shared causals under a bi-variate Gaussian distribution as 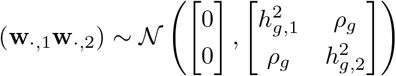 where 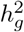 is the SNP-heritability for gene expression in tissue ·, and *ρ_g_* is genetic correlation. We repeated this for a total of 25 randomly sampled LD-blocks. We simulated complex trait for the GWAS panel as a linear combination of the genetic components of expression at causal genes. We first sampled causal genes at each LD-block with probability 1/*m_i_* where *m_i_* is the number of genes at block *i*. Then we computed 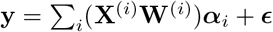 where 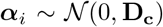 are causal effects for genes in the ithe LD-block, and 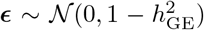. Next, we performed an association scan for (**y**, **X**^(1)^, …, **X**^(25)^) and computed SNP-trait Z-scores **z**_gwas_ using Wald statistics from linear regression. To perform a TWAS we fitted weights **Ω**^(1)^, …, **Ω**^(25)^ for the expression reference panel using GBLUP^36,37^ which were used to compute z_twas_. We then performed fine-mapping using the FOCUS algorithm on simulated z_twas_ vectors. Unless stated otherwise, simulation parameters were set to *N*_gwas_ = 50, 000, *N*_eQTL_ = 500, expression 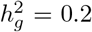 and trait 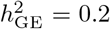 (i.e. variance explained in trait due to genetic component of gene expression^4^). For proxy-tissue simulations, we used values of proxy-tissue expression 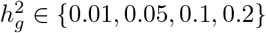 and *ρ_g_* ∈ {0, 0.1, 0.2, 0.3, 0.4, 0.5, 0.6, 0.7, 0.8, 0.9, 1.0}.

### Datasets

We downloaded publicly available summary statistics for lipids measurements GWAS.^17^ We filtered sites that were not bi-allelic, were ambiguous (i.e. allele 1 is reverse complement with allele 2), or had MAF less than 0.01. To perform TWAS on each of the lipids traits we used the software FUSION (see URLs). FUSION takes a summary-based approach to TWAS and requires as input GWAS summary statistics (i.e. SNP Z-scores) and eQTL weights. We downloaded publicly available expression weight data as part of the FUSION package. Reference LD was estimated in 1000 Genomes^40^ using 489 European individuals. Quality control, cis-heritability of expression, and model fitting have been described elsewhere.^1,4,33^ We prioritized adipose for our TWAS approach and used other reference panels as to act as proxy for adipose. That is, for all possible tissue-specific gene models in a region we first test predicted expression using adipose gene models. Then for the remaining genes found only in proxy tissue models, we select those with the best prediction accuracy (i.e. out-of-sample *R*^2^ normalized by complete-data 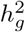 estimates). This resulted in 15,277 unique genes. Risk regions for FOCUS are ≈ 1Mb regions obtained from LDetect^28^ that contain at least one genome-wide significant SNP (*P*_gwas_ < 5 × 10^−8^).

## URLS

FOCUS: http://github.com/bogdanlab/focus/

FUSION: http://gusevlab.org/projects/fusion/

Lipids GWAS: http://lipidgenetics.org/

## Footnotes

1 Penalized regression may exhibit bias, but does so at the benefit of further reducing the mean-squared error, which is a measure of closeness to underlying parameters.

